# *Image matters*: Development of a novel staining method, the RGB-trichrome, as a reliable tool for the study of the musculoskeletal system

**DOI:** 10.1101/2020.03.18.996546

**Authors:** Francisco Gaytan, Concepción Morales, Carlos Reymundo, Manuel Tena-Sempere

## Abstract

Morphometry and histology are essential approaches for investigation and diagnosis of musculo-skeletal disorders. Despite the advent of revolutionary methods of image analysis and high resolution three-dimensional imaging technology, basic conventional light microscopy still provides an incisive overview of the structure and tissue dynamics of the musculoskeletal system. This is crucial to both preclinical and clinical research, since several clinically relevant processes, such as bone repair, osteoarthritis, and metabolic bone diseases, display distinct, if not pathognomonic, histological features. Due to the particular characteristics of the skeletal tissues (i.e., the existence of mineralized extracellular matrices), a large number of staining methods applicable to either decalcified or undecalcified tissues are available. However, it is usually the case that several staining methods need to be sequentially applied in order to achieve the different endpoints required to fully assess skeletal tissue structure and dynamics, and to allow morphometric quantification. We describe herein a novel staining method, the RGB trichrome, amenable for application to decalcified, paraffin embedded human musculoskeletal tissues. The acronym RGB corresponds to the three primary dyes used: picrosirius <u>R</u>ed, fast <u>G</u>reen, and alcian <u>B</u>lue. Although these individual pigments are commonly used either isolated, in binary combinations, or as part of more complex polychrome staining methods, when merged in the RGB trichrome staining produce high-quality/high-contrast images, permitting not only clear identification of different tissues (i.e., the different types of cartilage, bone and fibrous connective tissue), but also discrimination between calcified and uncalcified bone and cartilage, as well as an unexpected diversity of shades of color, while displaying singular properties among polychrome staining methods, such as the unveiling of the bone osteocyte dendritic/canalicular network. Hence, we propose the RGB trichrome as simple but highly-reliable tool for the preclinical and clinical study of the musculoskeletal system.

## Introduction

The musculoskeletal system provides mechanical support, permits movement and plays important endocrine and metabolic roles ^1^. It is composed of bone, cartilage and skeletal muscle, together with connecting structures, such as tendons and ligaments. The sustained increase in life expectancy witnessed in the last decades in low-, mid-, and high-income countries has resulted in an increase in the number and proportion of elderly people, and in the worldwide prevalence of age-related diseases ^2^. Aging of the musculoskeletal system is a leading cause of disability, and some of its age-related alterations, such as osteoporosis and osteoarthritis, are among the major epidemics of the 21st century, being, together with sarcopenia, highly-prevalent causes of mobility disability, and risk factors for falls and fractures in the elderly population ^3–7^.

Histology and histomorphometry are powerful tools for the study of skeletal structure and function, in both preclinical and clinical research ^8, 9^. Histopathological analysis of bone biopsies continues to be a diagnostic procedure and a research method in many skeletal diseases, such as osteoporosis, osteomalacia, hyperparathyroidism, renal osteodystrophy, Paget disease, and other metabolic bone diseases ^10^. Furthermore, the histological analysis is essential to assess different aspects of bone micro-architecture, the process of bone formation, mineralization, remodeling and fracture healing, as well as articular cartilage alterations ^11–13^. Due to the nature of the skeletal tissues, particularly the presence of a mineralized bone matrix, different tissue processing methods, either for decalcified or undecalcified tissues, have been developed. Paraffin-embedded tissues need to be previously decalcified, but resin-embedded undecalcified tissue processing is technically challenging, as it requires to be cut in special microtomes and the adaptation of conventional staining procedures ^14^.

A large number of staining methods are used for the recognition of different components and/or functional status of the musculoskeletal system. In order to highlight these different components and to allow histomorphometry quantification several staining methods are usually applied, from general staining, such as hematoxylin and eosin, Movat pentachrome, toluidine blue, alcian blue or safranin O/fas green ^9, 15^, to more specific silver impregnation ^16, 17^ or fluorescence-based ^18, 19^ methods for visualization of the osteocyte network. In addition, some specific staining methods, such as von Kossa or Masson-Goldner trichrome, are applied to undecalcified, resin-embedded, bone sections to distinguish the mineralized bone matrix from osteoid ^9^. However, a single general staining method highlighting the different tissue constituents, their interfaces and mineralized status, and the bone cell morphology, particularly as it pertains to the osteocyte dendritic/canalicular network, is not, to our knowledge, currently available.

In this study, we propose the use of a novel general staining method, the RGB trichrome; RGB being the acronym for the three primary color used: picrosirius <u>R</u>ed, fast <u>G</u>reen and alcian <u>B</u>lue. The sequential staining with these single dyes results in high-contrast staining, which allows the identification of the different components of the skeletal system, the clear differentiation of tissue interfaces and the visualization of associated fibrous and muscular tissues, permitting also to highlight the morphological characteristics of bone cells.

## Results

The RGB trichrome stain was applied to decalcified paraffin-embedded human tissues, including the different components of the skeletal system, such as cartilage and bone, as well as the associated skeletal muscle. Particular emphasis was made on the study of tissue interfaces, such as those between cartilage and bone, and calcified-uncalcified tissues.

### Cartilage

There are three types of cartilage, mostly depending on the different content of specific fibrous proteins in their matrices: hyaline, elastic and fibrocartilage, that are found in different locations in the human body.

Hyaline cartilage is the most abundant and shows two different types of extracellular matrix that, depending on the relative amount of ground substance (proteoglycans) and collagen, displayed a differential staining when the RGB trichrome was applied. Accordingly, the pericellular territorial matrix (with high proteoglycan content) showed blue-purple staining, whereas the interterritorial matrix (containing less proteoglycans and more collagen) presented an intense red staining (**Figs. 1A,B**). Inside the lacunae, chondrocytes were blue/green stained. Although the structural arrangement of elastic cartilage is similar to that of hyaline cartilage, its staining properties with the RGB trichrome were rather different, and showed blue/green stained intercellular matrix (**Figs. 1C,D**). Otherwise, fibrocartilage associated to the attachment of tendons or ligaments to bone (i.e., enthesis region) displayed round chondrocytes dispersed in pairs or small groups in longitudinal rows, showing a blue-stained pericellular matrix, interposed between dense, red-stained, collagen bundles in uncalcified fibrocartilage or in a beige/greenish-stained background in calcified fibrocartilage (**Figs. 1 E,F**).

**Figure 1.**
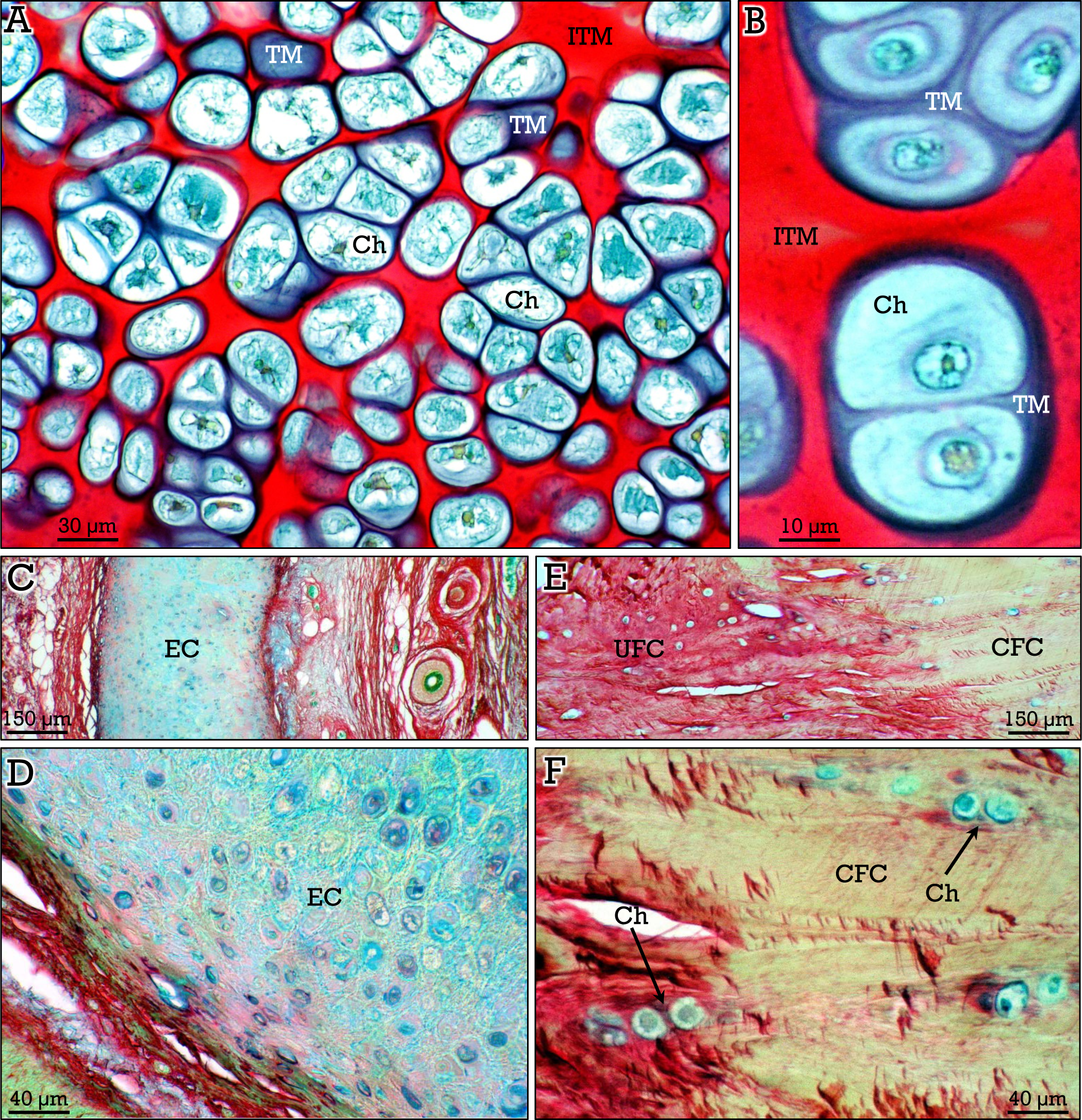
RGB trichrome staining of cartilage. A,B, hyaline cartilage shows chondrocytes (*Ch*) surrounded by blue/purple stained territorial matrix (*TM*) and red-stained interterritorial matrix (*ITM*). C,D, elastic cartilage (*EC*) showing blue/green stained matrix. E,F, chondrocytes (*Ch*) are present in low number in uncalcified (*UFC*, red-stained) and calcified (*CFC*, beige/greenish stained) fibrocartilage.

### Cartilage-bone interface

Cartilage-bone interfaces play essential roles in the maintenance of structural and functional integrity of different skeletal structures, such as the growth plate during endochondral ossification, diarthrodial joints (i.e., articular cartilage-subchondral bone interface), or during skeletal tissue repair ^20^. Osteochondromas are the most frequent benign bone tumors ^21^, and usually arise at the metaphysis of long bones. They are composed of a cap of proliferating hyaline cartilage undergoing endochondral ossification with subjacent cancellous bone formation, surrounding marrow spaces continuous with the marrow of underlying bones. They appear as developmental lesions (rather than true tumors) resulting from separation and outgrowth of the growth plate. Osteochondromas constitute, for the purposes of this study, an excellent model to analyze bone-cartilage interface in humans, in which normal tissues, such as growth plates, are not usually present.

Young (growing) osteochondromas resembled the epiphyseal growth plate, showing close similar histologic features ^11, 22^. Hypertrophic chondrocytes were present and a clear contrast was found between red-stained cartilage matrix and green-stained bone matrix (**Figs. 2A**). In the areas facing subchondral bone, pink/greenish-stained calcified cartilage matrix was invaded by connective-vascular tissue (**Figs. 2 A,B**). Bone trabeculae showed peripheral zones of bone matrix and central zones of calcified cartilage matrix, together with occasional residual blue-stained chondrocyte lacunae (**Figs. 2B,C**). Cement lines between cartilage and bone matrix were evident (**Fig. 2C**). In some deep invagination areas, resorption of cartilage by osteo(chondro)clasts was observed (**Fig. 2D**).

**Figure 2.**
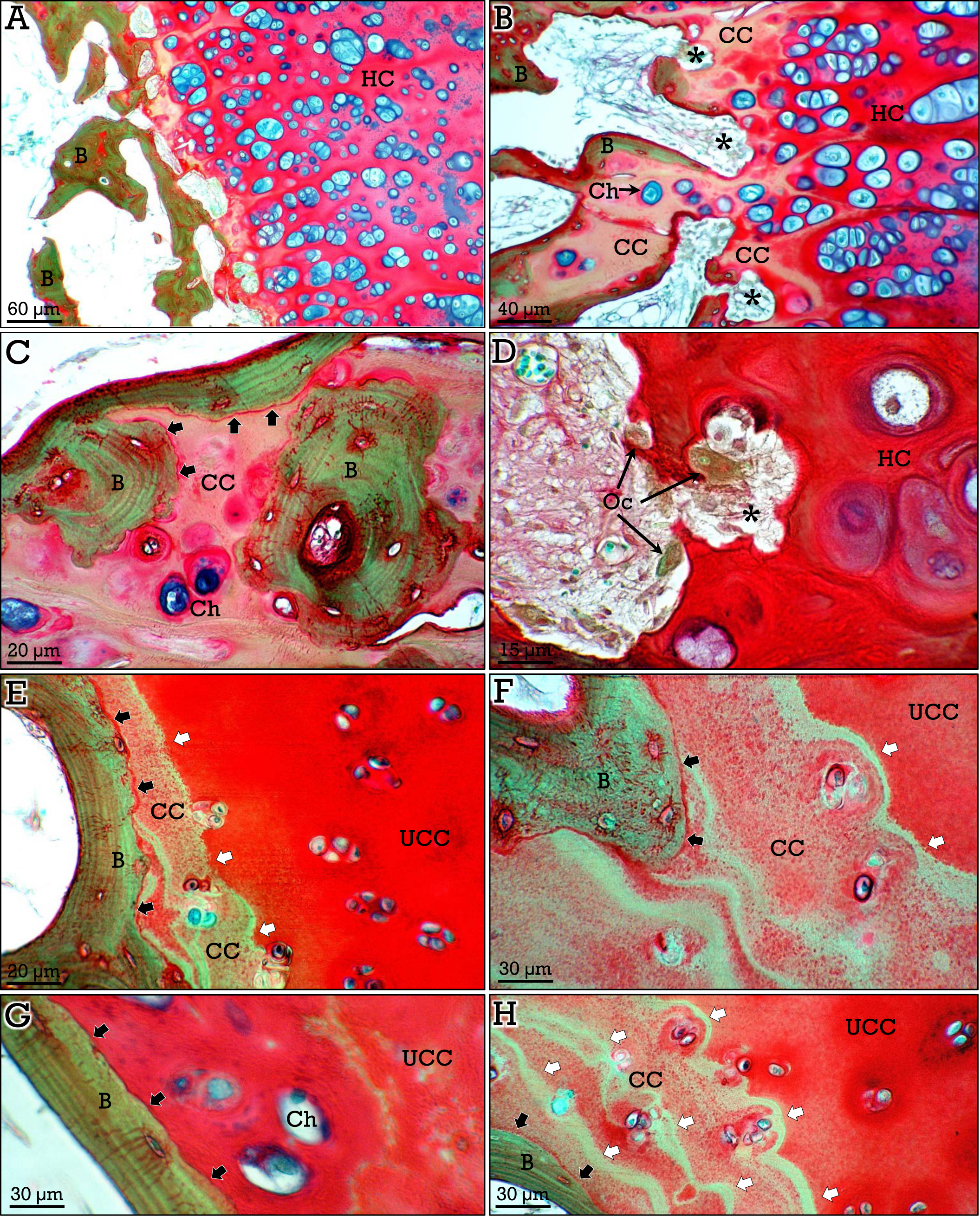
RGB trichrome staining of the cartilage-bone interface in osteochondromas. In growing osteochondromas (**A-D**), resembling the epiphyseal growth plate, the hyaline cartilage (*HC*) is undergoing endochondral ossification, showing calcified matrix (*CC*) at the bone interface, invasion by connective and vascular tissue (*asterisks*) and formation of bone trabeculae (*B*) containing a center of calcified cartilage, and occasional chondrocytes (*Ch*). The cement lines between bone and calcified cartilage can be observed (*black arrows* in **C**). Osteo(chondro)clasts (*Oc*) can be observed in the resorption zones. In non-growing osteochondromas (**E-H**), resembling articular cartilage, chondrocytes were sparse and non-hypertrophic, and a tidemark (*white arrows*) can be observed between uncalcified (*UCC*) and calcified (*CC*) cartilage matrix, as well as cement lines (*black arrows*) between cartilage and bone. In some areas, a direct contact between uncalcified cartilage and bone (*black arrows* in **G**) or reduplication of the tidemark (*white arrows* in **H**) can be observed.

Otherwise, mature (non-growing) osteochondromas resemble articular cartilage ^11^. Chondrocytes were sparse and not hypertrophic, and a clear tidemark was observed separating non-calcified (red-stained) from calcified (pink/greenish-stained) cartilage matrix (**Figs. 2E,F**), equivalent to the tidemark present in articular cartilage ^23, 24^. Cement lines separating cartilage and bone matrix were also evident (**Figs. 2E,F**). However, a direct contact between uncalcified cartilage and bone was also observed (**Fig. 2G**), similar to that reported in some places in articular cartilage ^25^. Furthermore, reduplication of the tidemark was observed in some areas of the cartilage-bone interface (Fig. 2H), resembling the features of articular cartilage in osteoarthritis ^24^.

### Bone

Woven and lamellar bone showed similar green-stained matrix. Woven bone, characterized by the irregular arrangement of collagen fibers and osteocytes, was abundant in bone-forming osteoid osteomas that will be described in later sections. Otherwise, the two types of lamellar bone, compact and cancellous, showed the same staining properties. A very relevant aspect of the RGB trichrome staining method, as compared to other available trichrome staining protocols, was the unveiling of the osteocyte dendritic/canalicular network (**Fig. 3**). Osteocyte dendrites/canaliculi appeared red-stained and could be clearly appreciated, even in canalicular cross-sections, contrasting against the green-stained bone matrix (**Fig. 3**). In compact bone, osteocytes were arranged circumferentially around the central vascular channels in osteons (**Fig. 3A,B**). In cancellous bone, Haversian systems were usually not found and osteocytes were in general parallel to the surface of the trabeculae (**Fig. 3C,D**). Differences in the density of osteocytes and canalicular network were observed (**Fig. 3E**), and the nucleus and cytoplasm of the osteocytes were clearly observed inside the lacunae (**Fig. 3F,G**).

**Figure 3.**
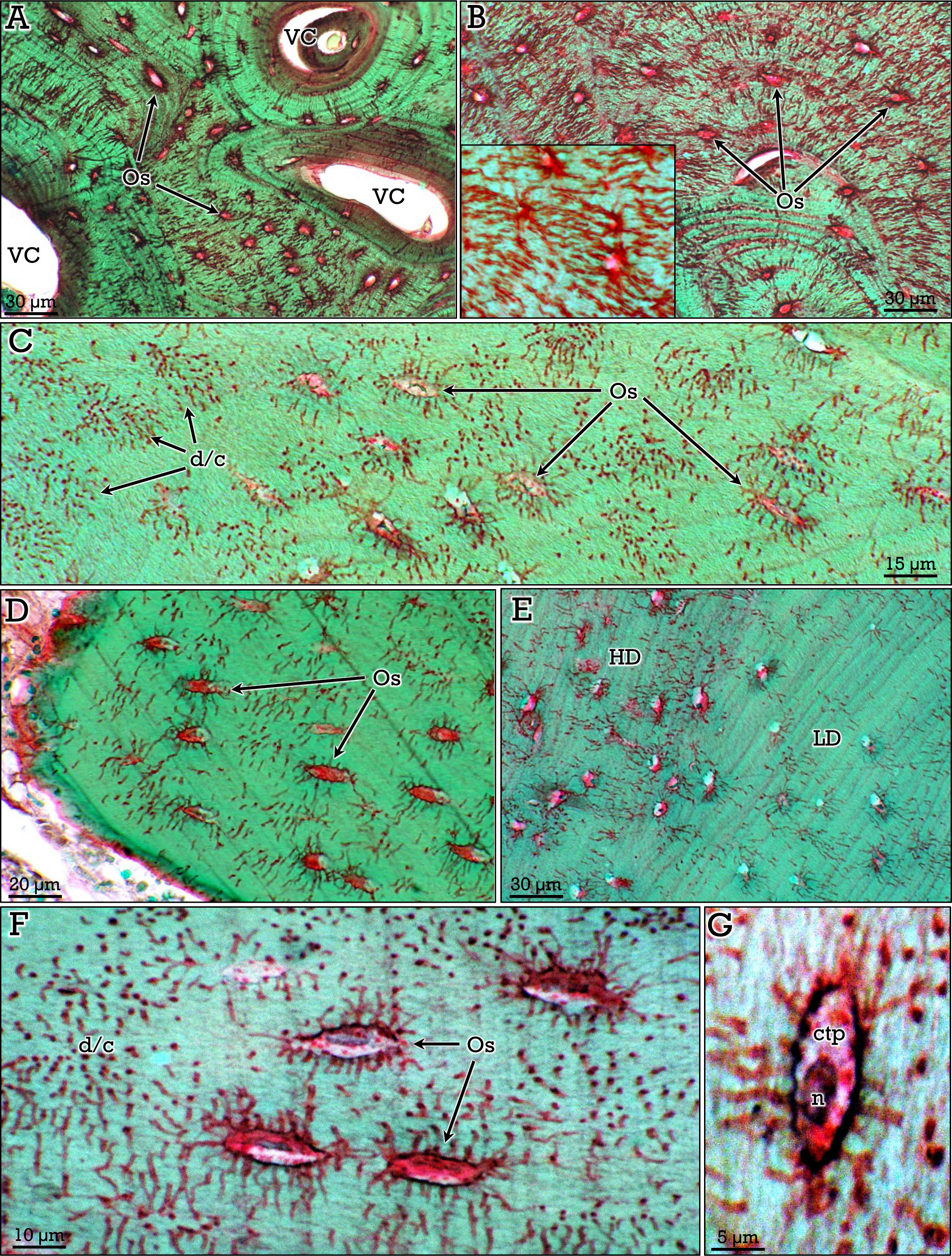
RGB trichrome staining of bone. In compact bone (**A, B**), osteocytes (*Os*) are arranged concentrically around vascular channels (*VC*), showing abundant dendrites/canaliculi. In cancellous bone (**C**-**G**), osteocytes are arranged in parallel with respect to the surface of the trabeculae, distributed in areas of high (*HD*) and low (*LD*) densities. Osteocyte dendrites/canaliculi (*d/c* in **C** and **F**), as well as the nuclei (*n*) and cytoplasm (*ctp*) of the osteocytes (**F,G**) can be observed.

A particularly important aspect of bone histology is the distinction between unmineralized (i.e., osteoid) and mineralized bone matrix. The presence of an increased amount of osteoid occurs in several bone alterations involving defective matrix mineralization ^9^. In RGB-trichrome stained bones, osteoid appears as red-stained seams in the surface of some trabeculae (**Fig. 4A**), showing a high contrast with mineralized bone (**Fig. 4**). Osteoid was very abundant in osteoid osteomas (a benign bone-forming tumors; see **Fig. 5**) that lay down bone matrix rapidly and show a mixture of uncalcified (**Fig. 5A**), partially calcified (**Fig. 5B**), and calcified (**Fig.5C**) trabeculae. In these tumors, abundant fibrous connective tissue was present in the marrow spaces, as well as traversing the newly formed bone matrix (**Figs. 5E**). Calcified bone corresponded to woven bone with irregular distribution of osteocytes that show few and short dendrites/canaliculi (**Figs. 5F**).

**Figure 4.**
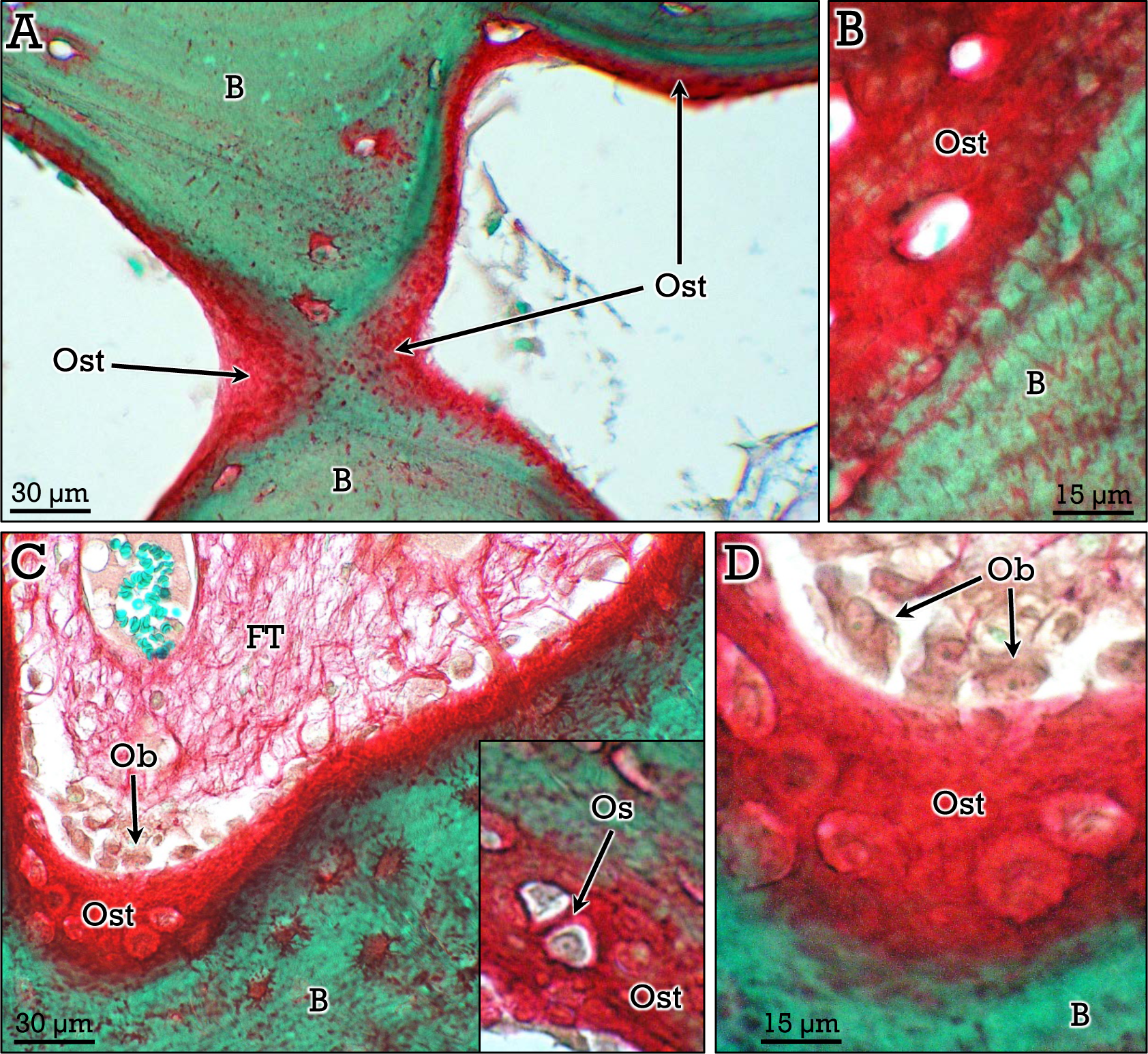
Differential staining of osteoid and bone with the RGB trichrome method. Osteoid seams (*Ost*) at the surface of bone (*B*) trabeculae (see **A,B**), and osteoid formation by osteoblasts (*Ob* in **C,D**), that become trapped and differentiate to osteocytes (*Os* in the inset in **C**). *FT*, fibrous tissue in the marrow.

**Figure 5.**
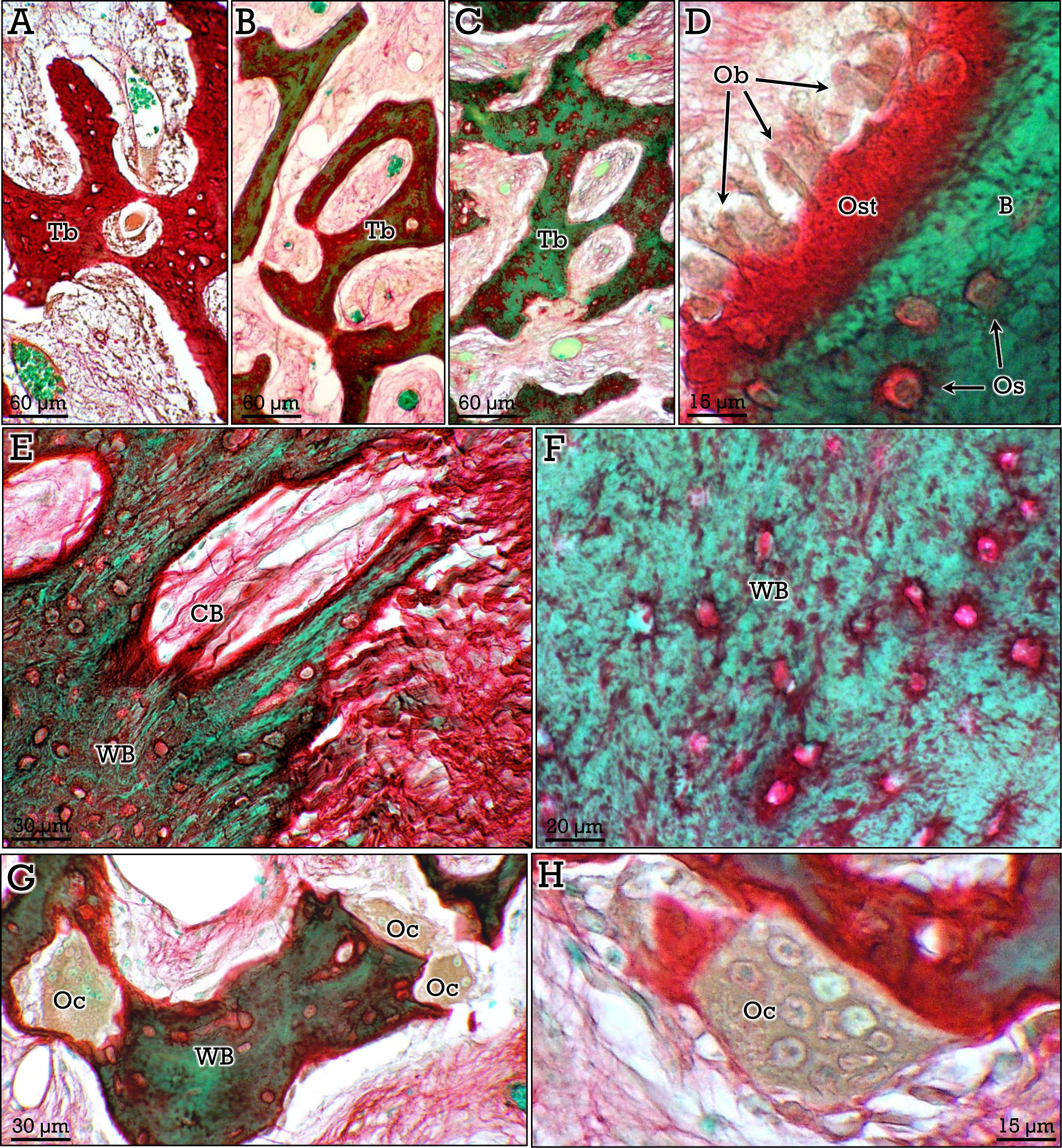
Staining of osteoid osteomas with the RGB staining method, showing bone formation and resorption. Trabeculae (*Tb*) show different degrees of calcification, from osteoid (**A**), to partially calcified (**B**) and calcified (**C**) matrix. The surface of the trabeculae show abundant and prominent osteoblasts (*Ob* in **D**). Osteoblasts (*arrows* in **D**) lay down bone matrix in the scaffold of dense fibrous tissue. Newly formed woven bone (WB, in **E, F**) is crossed by collagen fiber bundles (CFB), which are evident even after partial calcification (*CB* in **E**). Multinucleated osteoclasts (*Oc*) in the surface of trabeculae of woven bone (**G, H**) are also observed.

Osteoblasts appear as large cells lining the osteoid surface (see **Fig. 4C,D**), and were very abundant in osteoid osteomas (**Fig. 5D**). After being trapped in the osteoid matrix, they differentiate into osteocytes (**Figs. 4C** & **5D**). The third specific bone cell type, namely, osteoclasts, were also abundant in bone forming tumors. They appeared as multinucleate cells located in resorption lacunae, in either osteoid or woven bone tissue (**Fig. 5G,H**).

### Skeletal muscle

Skeletal muscle is closely associated to skeleton and is structurally and functionally linked to bones and joints. Skeletal muscle stained with the RGB trichrome showed pale green-stained muscle fibers surrounded by the red-stained collagen in the covering connective tissue (**Fig. 6A**). The striated nature of muscle fibers can be easily appreciated in longitudinal sections (**Fig. 6B**). The RGB trichrome staining also provides a high contrast between muscle fibers and collagen at myo-tendinous junctions (**Fig. 6C**).

**Figure 6.**
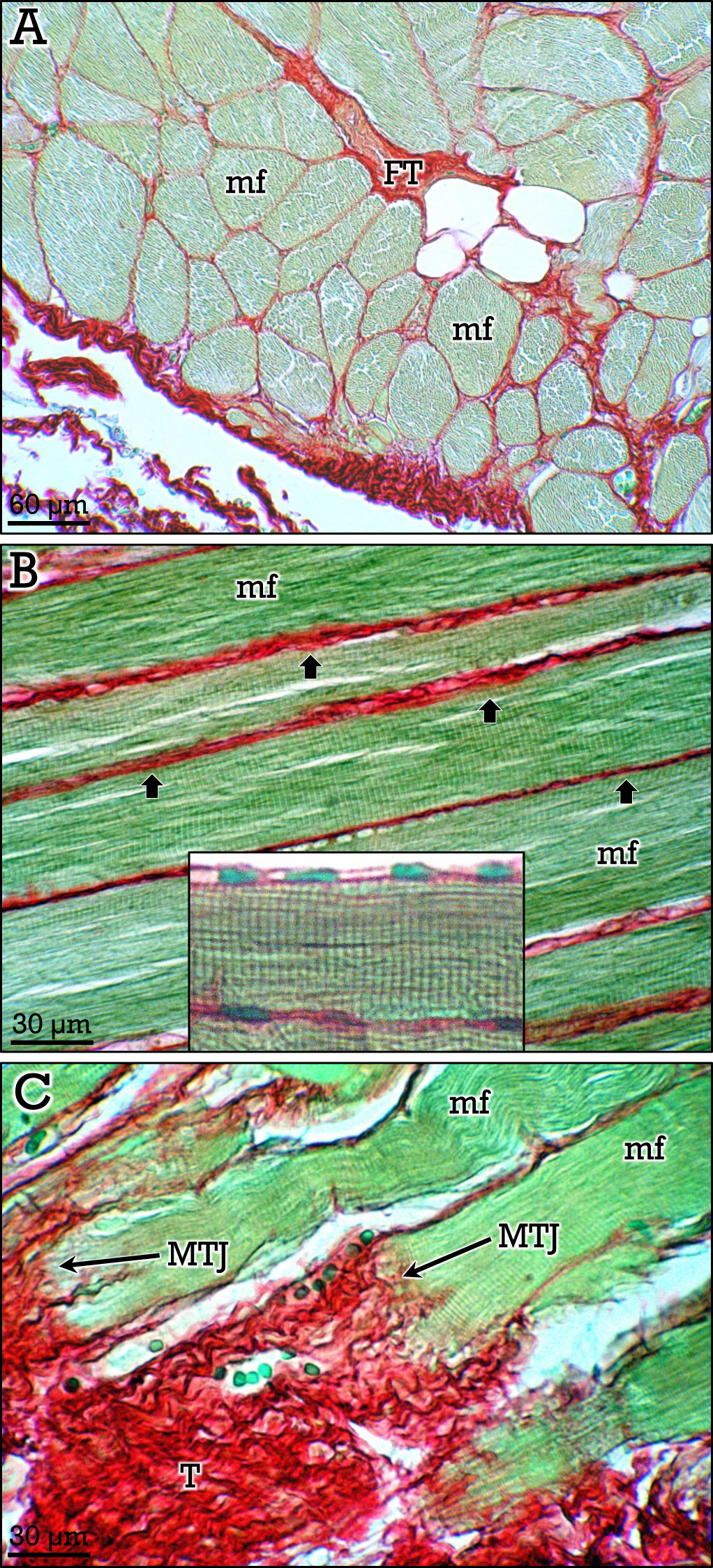
RGB trichrome staining of skeletal muscle. In **A**, muscular fibers (*mf*) and fibrous tissue (*FT*) can be identified. In longitudinal sections (**B**), the endomisium (*arrows*) and the cytoplasmic striations and subsarcolemal nuclei (*inset*) can be observed. In **C**, differential staining of the muscle fibers (*mf*) and fibrous tissue (*FT*) at myo-tendinous junction (*MTJ*).

### Musculoskeletal system during embryonic development

The RGB trichrome staining was also applied to first trimester embryos to analyze its potential as a topographic staining for embryonic tissues, and particularly for the components of the developing musculoskeletal system. Embryonic hyaline cartilaginous structures showed scarce interterritorial matrix and high density of chondrocytes (**Figs. 7A,B**). A clear differential staining was found between embryonic connective tissue (blue-stained), collagenous tissue (red-stained), myotubes (green-stained), and hyaline cartilage (**Figs. 7C-F**). Early intramembranous ossification was also clearly identified (**Fig. 7B**).

**Figure 7.**
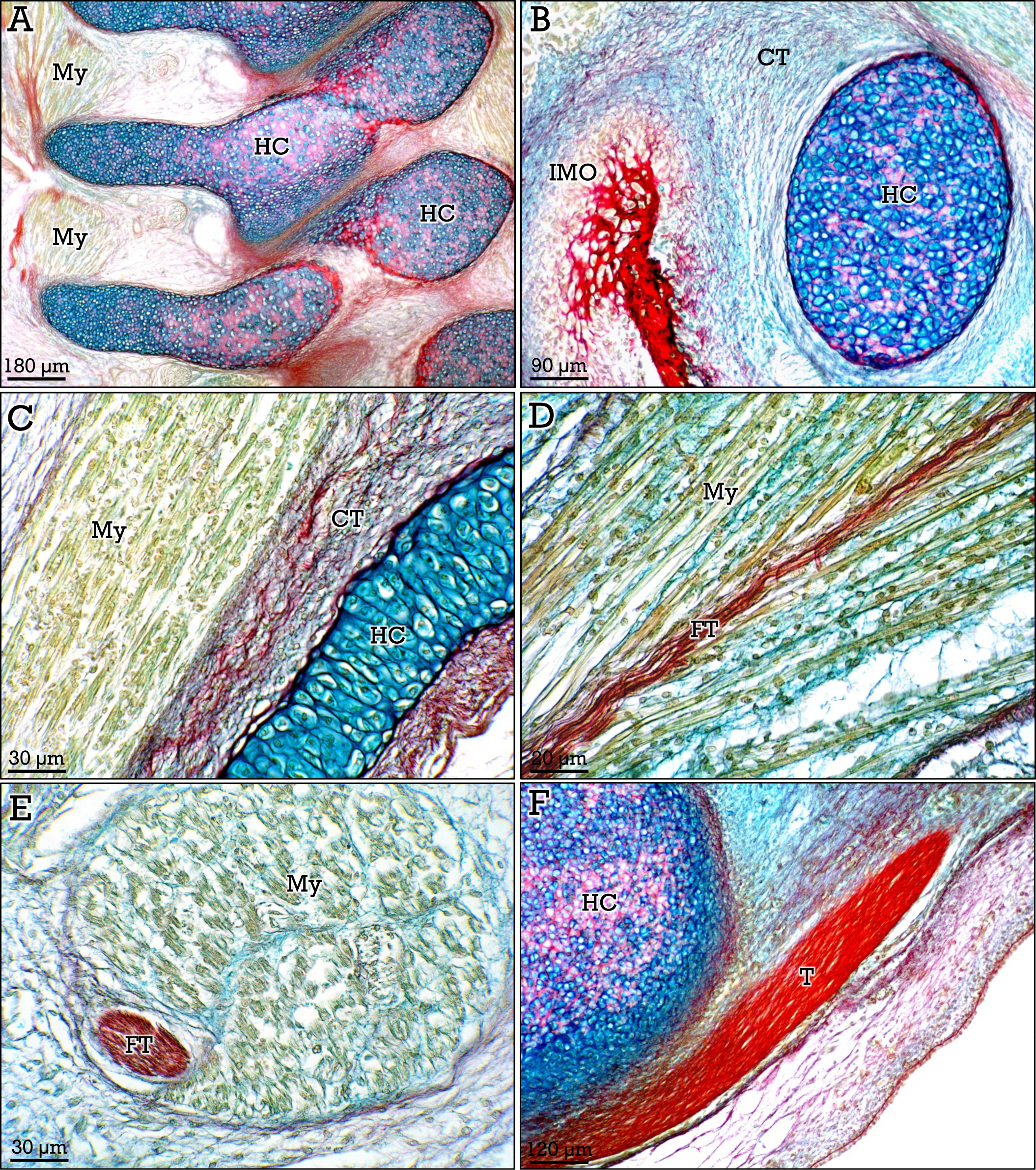
Staining of embryonic musculoskeletal tissues with RGB trichrome. Hyaline cartilage forming cartilage anlages (*HC*) show a high density of chondrocytes and scarce interterritorial matrix (see **A,B**). Myotubes (*My*) are clearly differentiated from connective tissue (*CT* in **C**). **D,E**, myotubes (*My*) and dense fibrous tissue (*FT*) in longitudinal (**D**) and transverse (**E**) embryonic muscle sections. In **F**, insertion of a tendon (*T*) in the cartilaginous anlage. Initial intramembranous ossification (*IMO* in **B**) is also observed.

### Cartilage pathological samples

Although staining of overtly pathological tissue samples was beyond the purposes of this study, we included some sections of chondrosarcomas, a malignant bone tumor that produce chondroid (cartilaginous) matrix, to reveal the unexpected spectrum of shades of color provided by RGB trichrome staining method, due to the heterogeneity of tumor cells and extracellular matrix (**Fig. 8**).

**Figure 8.**
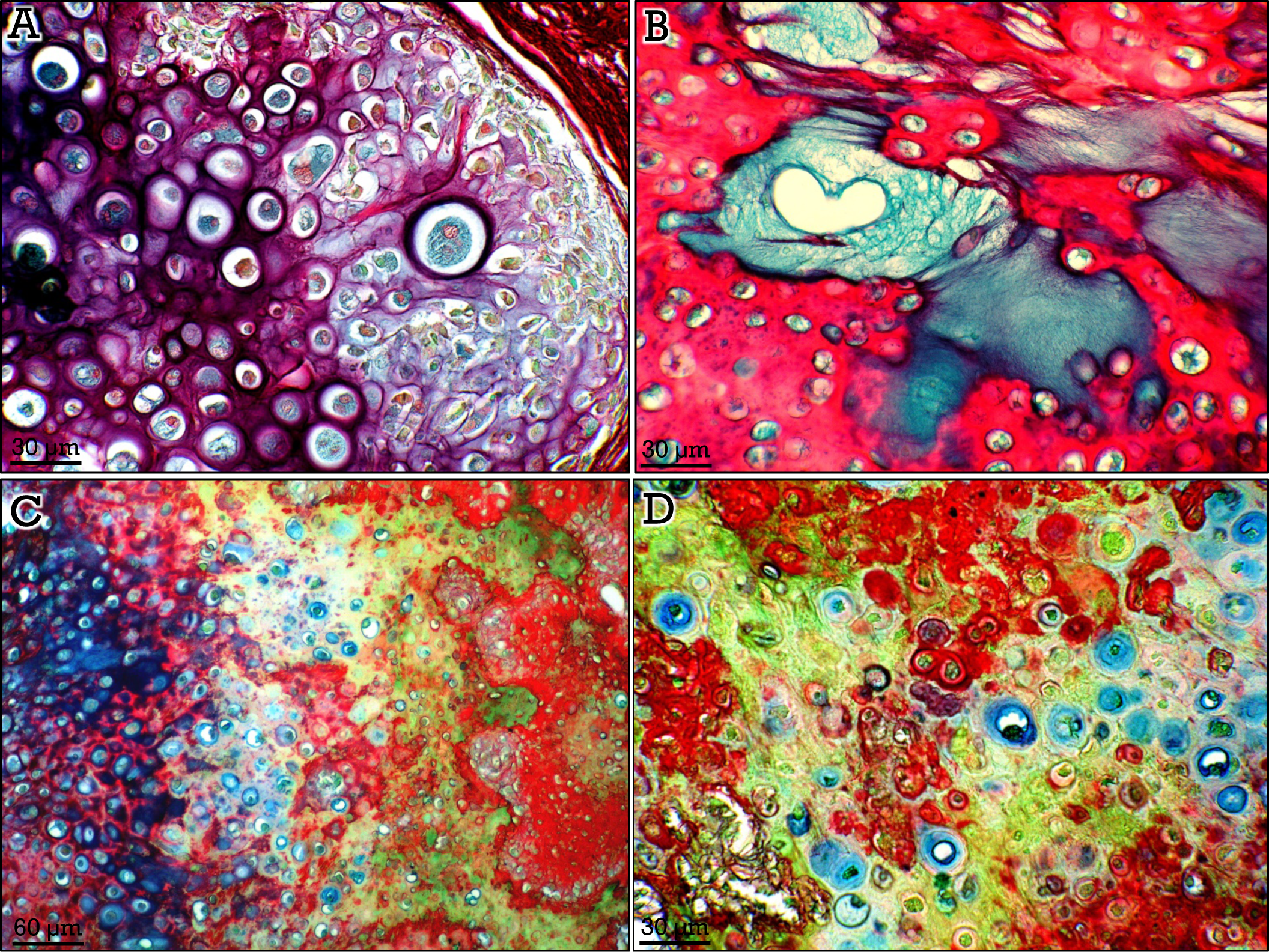
RGB trichrome staining of chondrosarcomas. RGB trichrome staining of sections from two different chondrosarcomas (**A,B** vs. **C,D**), showing a different and heterogeneous staining and a high variety of shades of colors.

## Discussion

The development of new techniques, such as confocal and third harmonic generation microscopy, as well as scanning electron microscopy-associated techniques, has provided a new vision of the skeletal tissue micro-architecture ^19, 26–29^. However, these microscopy technologies remain expensive, require highly-specialized management, and are not available in many laboratories. In the field of light microscopy, numerous staining methods have been developed, and usually several staining procedures are needed to cope with the differences in tissue components and functional status of the skeletal system ^9, 15^. In this context, a general staining method should fulfill a number of requirements to provide a general overview of the microstructure and functional characteristics of the musculo-skeletal organs, at cell and tissue levels. These requirements should include (i) a clear distinction among cartilage, fibrous collagenous tissue and bone, (ii) a clear differentiation between uncalcified and calcified tissues (either cartilage, bone or tendon), (iii) the identification of general bone structure (i.e., woven vs. lamellar bone), (iv) the discrimination of intramembranous bone formation vs. endochondral ossification, and (v) the identification and characterization of bone cells (i.e., osteoblasts, osteoclasts, and particularly, the osteocyte lacuno-canalicular network). Whereas all these endpoints can be achieved by using a combination of different staining methods, a single general staining fulfilling all these requirements was, to our knowledge, not available as yet.

The RGB trichrome is a simple staining method that provides a clear tissue discrimination through distinctive coloring of the main components of the musculoskeletal system. Thus, hyaline cartilage appears blue/red (territorial/interterritorial matrix), mineralized cartilage matrix appears pink/greenish, osteoid appears red, and mineralized bone appears green. In addition, the fronts delimiting the high contrasted tissue interfaces (i.e., tidemarks and cement lines) can be clearly identified. In addition, skeletal muscle and collagen (in muscles and myo-tendinous junctions) are differentially stained and even the thin layer of fine red-stained collagen fibers (endomysium) surrounding each individual (green-stained) muscle fiber can be appreciated. Staining of different types of cartilage (i.e., hyaline vs. elastic) results in differential matrix staining, thus showing the sensitivity of this staining method to relatively small differences in the composition of the extracellular matrix. An extreme example of this sensitivity is provided by the staining of chondrosarcomas that results in a wide spectrum of shades of colors in the tumoral intercellular matrix. These color differences are a reflection of the heterogeneity of tumor extracellular matrix composition, are expected to occur also among different tumors, and could be related to differences in tumor biology and behavior.

Identification and characterization of bone cells are also important to analyze bone micro-architecture and dynamics. Early studies on the process and regulation of bone remodeling were mainly focused in the bone cell types acting as final effectors of bone resorption and synthesis, namely, osteoclasts and osteoblasts, respectively. However, in the last decades, the osteocytes, that constitute about 90% of bone cells and are long-lived, have been pointed out as the most important regulators of bone dynamics, acting as mechano-sensors ^30^, as well as endocrine-like cells releasing signaling molecules and regulatory factors, such as fibroblast growth factor 23 (FGF23), which target other cell types and organs ^31, 32^. Indeed, a recent study raised the intriguing possibility that osteocytes participate in the regulation of fat body mass through sensing changes in bone strain ^33^. In this context, osteocytes seem to play central roles in bone physiology and pathophysiology ^32, 34–37^; osteocyte dendritic network being the morphological counterpart of the functional coupling of osteocytes among them and with other bone cell types ^26, 29, 30^. Although ground bone tissue sections and classical staining methods, such as silver impregnation ^17, 38^, allow visualization of the osteocyte lacuna-canalicular network, the novel high-resolution and 3D imaging technologies have provided during the last decade detailed views of the osteocyte syncytia ^26–29, 39, 40^. Notwithstanding, the high technical demands of such approaches hamper their universal implementation. The RGB trichrome proposed herein is a simple staining method that does not require special technology but provides a high-quality first approach to the analysis of the osteocyte dendritic/canalicular network, as it permits to discern the general distribution and density of osteocytes, as well as their dendritic processes. This, together with the tri-chromic discrimination between different tissue types, constitutes one of the most relevant outcomes of this novel staining.

Furthermore, the RGB trichrome seems to be an excellent topographical staining method for embryos, especially for mapping the development of the different components of the musculoskeletal system. Indeed, by applying the RGB trichrome, the developing muscle cells (myotubes), dense fibrous tissue in developing tendons, hyaline cartilage and intramembranous ossification centers can be easily identified.

In sum, we propose herein a novel, as yet simple staining method, the RGB trichrome, which constitutes an undemanding and reliable procedure to visualize the micro-structure, and indirectly assess the functional status, of the different components of the musculoskeletal system and their tissue interfaces, while unveiling the osteocyte dendritic/canalicular system. Considering its high-quality output (including beautiful and color-vivid images) and its minimal technical demands, the RGB trichrome will likely become the first-step choice for the histopathological analysis of the musculo-skeletal system, in clinical and experimental research.

## Materials and Methods

### Human musculoskeletal tissues

Human tissues were obtained from the Bio-archive of the Department of Pathology of the Faculty of Medicine and Nursing of the University of Cordoba. These specimens were collected for diagnostic purposes between 1980 and 2000, in keeping with contemporary legislation and standard clinical practices, which included informed consent and general Institutional Board revision. The use of the samples was authorized by responsible members of the Department of Pathology, who granted strict adherence to current legislation regarding patient confidentiality.

Paraffin-embedded tissue samples, corresponding to 2 embryos (∼9 week) from spontaneous abortions, 6 bone biopsies, 3 osteochondromas, 2 osteoid osteomas, 2 chondrosarcomas, 4 muscle biopsies, and 2 samples from external ears, were used in this study, after initial classification by members of the Department of Pathology, and re-evaluation by the co-authors, CM and CR, as experienced pathologists. Tissues had been fixed in neutral buffered formaldehyde, decalcified in 5% nitric acid, and routinely embedded in paraffin. Six micrometer-thick sections were cut and, after dewaxing and rehydration, used for RGB trichrome staining.

### Reagents

For the RGB trichrome staining method, the following reagents were used: Sirius Red (also termed Direct Red 80; Ref# 365548); Fast Green FCF (Ref#F7258); and Alcian Blue 8GX (Ref#A5268). All reagents were obtained from Sigma Aldrich (St. Louis, MO).

### Staining protocol

Sections were stained sequentially with 1% alcian blue 8GX in 3% aqueous acetic acid solution (pH 2.5) for 20 min and rinsed in tap water for 5 min. Thereafter, the sections were stained with 1% fast green FCF in distilled water for 20 min and rinsed in tap water for 5 min. Finally, the sections were stained with 1% sirius red in saturated aqueous solution of picric acid (i.e., picrosirius red) for 30 min, rinsed in two changes of 5 min in acidified water (1% acetic acid in tap water), dehydrated in two changes of 100% ethanol, cleared in xylene, and mounted in a resinous medium.

## Acknowledgement

The authors acknowledge the superb technical assistance of Mr. Esteban Tarradas, responsible of the Scientific Image Unit at the Faculty of Medicine of the University of Cordoba, in the preparation of the figures of this manuscript.

## Author Contributions

F.G. had a leading role in the design of the novel staining method and the implementation of all tissue analyses, with the close assistance of C.M. and C.R., expert pathologists, which gave access also to the archival tissue samples used in this study. M.T.S. collaborated in data analysis and interpretation/discussion of the results. F.G. was responsible of preparation of the first draft of the manuscript, which was revised and edited by M.T.S. All authors revised and approved the paper.

